# Opposing modulation of Cx26 gap junctions and hemichannels by CO_2_

**DOI:** 10.1101/584722

**Authors:** Sarbjit Nijjar, Daniel Maddison, Louise Meigh, Elizabeth de Wolf, Thomas Rodgers, Martin Cann, Nicholas Dale

## Abstract

Cx26 hemichannels open in response to moderate elevations of CO_2_ (PCO_2_ 55 mmHg) via a carbamylation reaction that depends on residues K125 and R104. Here we investigate the action of CO_2_ on Cx26 gap junctions. Using a dye transfer assay, we found that an elevated PCO_2_ of 55 mmHg greatly delayed the permeation of a fluorescent glucose analogue (NBDG) between HeLa cells coupled by Cx26 gap junctions. However, the mutations K125R or R104A abolished this effect of CO_2_. Whole cell recordings demonstrated that elevated CO_2_ reduced the Cx26 gap junction conductance (median reduction 5.6 nS, 95% confidence interval, 3.2 to 11.9 nS) but had no effect on Cx26^K125R^ or Cx31 gap junctions. CO_2_ can cause intracellular acidification, but using 30 mM propionate we found that acidification in the absence of a change in PCO_2_ caused a median reduction in the gap junction conductance of 5.3 nS (2.8 to 8.3 nS). This effect of propionate was unaffected by the K125R mutation (median reduction 7.7 nS, 4.1 to 11.0 nS). pH-dependent and CO_2_-dependent closure of the gap junction are thus mechanistically independent. Mutations of Cx26 associated with the Keratitis Ichthyosis Deafness syndrome (N14K, A40V and A88V) also abolished the CO_2_-dependent gap junction closure. Elastic network modelling suggests that the lowest entropy state when CO_2_ is bound, is the closed configuration for the gap junction but the open state for the hemichannel. The opposing actions of CO_2_ on Cx26 gap junctions and hemichannels thus depend on the same residues and presumed carbamylation reaction.

## Introduction

The canonical function of connexins is to form intercellular junctions between cells –gap junctions – through the docking of hexameric connexons in the opposing membrane of each cell. However, the individual connexons –known as hemichannels –can also function on their own (Stout *et al.*, 2004; Weissman *et al.*, 2004; Pearson *et al.*, 2005; Huckstepp *et al.*, 2010b). Hemichannels act as plasma membrane channels, which in addition to mediating transmembrane ionic currents, also permit the transmembrane fluxes of small molecules such as ATP (Stout *et al.*, 2002; Pearson *et al.*, 2005; Kang *et al.*, 2008; Huckstepp *et al.*, 2010a).

We have studied the hemichannels of connexin26 (Cx26) and found that these hemichannels are directly gated by CO_2_ (Huckstepp *et al.*, 2010a; Meigh *et al.*, 2013; Meigh *et al.*, 2014). CO_2_ opens the hemichannel and permits the efflux of ATP, which can act as a neurotransmitter. This is particularly important in the CO_2_ sensitive control of breathing where Cx26 hemichannels in the medulla oblongata act as novel chemosensory transducers (Huckstepp *et al.*, 2010a; Huckstepp & Dale, 2011). Cx26 hemichannels also impart CO_2_ sensitivity to dopaminergic neurons of the substantia nigra and GABAergic neurons of the ventral tegmental area (Hill *et al.*, 2020). The action of CO_2_ on Cx26 has been proposed to occur via carbamylation of the residue K125. The carbamylated lysine forms a salt bridge to R104 of the neighbouring subunit. These intersubunit carbamate bridges bias the hemichannel to the open configuration (Meigh *et al.*, 2013). Extensive evidence supports a direct action of CO_2_ on the hemichannel rather than an indirect effect via pH: 1) in isolated inside-out or outside-out patches changes in PCO_2_ at constant pH alter Cx26-gating (Huckstepp *et al.*, 2010a); 2) insertion of the carbamylation motif into Cx31, which is insensitive to CO_2_, creates mutant Cx31 hemichannels that can be opened by CO_2_ (Meigh *et al.*, 2013); 3) mutation of the key residues K125 and R104 to respectively arginine and alanine destroys CO_2_-sensitivity of the hemichannel; 4) the mutations K125E and R104E (in effect engineering the action of CO_2_ into the subunit via the carboxy group of glutamate) create constitutively open hemichannels which are CO_2_-insensitive (Meigh *et al.*, 2013); 5) the mutation K125C creates a hemichannel that can be opened with NO or NO_2_ via a nitrosylation reaction on the cysteine residue at 125 and subsequent salt bridge formation to R104 (Meigh *et al.*, 2015); and 6) the double mutation K125C and R104C creates a redox-sensitive hemichannel presumably via disulfide bridge formation between the cysteine residues at 104 and 125 (Meigh *et al.*, 2015).

Although the actions of CO_2_ on Cx26 hemichannels are well characterized, we have not studied the action of CO_2_ on Cx26 gap junctions. When the two connexons dock to form a complete gap junction, there is likely to be significant conformational rearrangement and constraint of the resulting dodecameric complex. Therefore, we cannot assume that CO_2_ will modulate a complete gap junction in the same way as a hemichannel. There has been a previous study on the closing effect of CO_2_ on Cx32 and Cx26 gap junctions expressed in *Xenopus* oocytes (Young & Peracchia, 2004). This prior study used non-physiological conditions in which both the extracellular and intracellular pH would become very acidic: exposure to 30-100% CO_2_ in the absence of bicarbonate in the extracellular medium. This study most likely reported an effect of pH on the gap junction. In this paper, we report the actions of much lower physiological concentrations of CO_2_ (~9%), in a CO_2_/HCO_3_^−^ buffered system at constant extracellular pH, on gap junctions formed between pairs of HeLa cells expressing Cx26. We find that modest increases in PCO_2_ *close* complete gap junctions, and that this is most likely a direct effect mediated through its binding to the same residues that result in the *opening* of the hemichannel. This result reinforces the need to develop further high-resolution structures for Cx26 hemichannels and gap junctions with and without CO_2_ bound.

## Methods

### HeLa cell culture and transfection

HeLa DH (ECACC) cells were grown in DMEM supplemented with 10% fetal bovine serum, 50 μg/mL penicillin/streptomycin and 3 mM CaCl_2_. For electrophysiology and intercellular dye transfer experiments, cells were seeded onto coverslips in 6 well plates at a density of 2×10^4^ cells per well. After 24 hrs the cells were transiently transfected with Cx26 constructs tagged at the C-terminus with a fluorescent marker (mCherry) according to the GeneJuice Transfection Reagent protocol (Merck Millipore). The HeLa cells stably expressing mouse Cx26 were originally obtained from Dr K. Willecke (Elfgang *et al.*, 1995).

### Cx26 mutants

The mutations used in this study were introduced into the Cx26 gene by QuikChange site directed mutagenesis and have been described previously (Meigh *et al.*, 2013) (Cook *et al.*, 2019).

### Solutions used

#### Standard aCSF

124 mM NaCl, 3 mM KCl, 2 mM CaCl_2_, 26 mM NaHCO_3_, 1.25 mM NaH_2_PO_4_, 1 mM MgSO_4_, 10 mM D-glucose saturated with 95% O2/5% CO_2_, pH 7.5, PCO_2_ 35 mmHg.

#### Hypercapnic aCSF

100 mM NaCl, 3 mM KCl, 2 mM CaCl_2_, 50 mM NaHCO_3_, 1.25 mM NaH_2_PO_4_, 1 mM MgSO_4_, 10 mM D-glucose, saturated with 9% CO_2_ (with the balance being O_2_) to give a pH of 7.5 and a PCO_2_ of 55 mmHg respectively.

#### Propionate solution

82mM NaCl, 30 mM Na-propionate, 3 mM KCl, 2 mM CaCl_2_, 26 mM NaHCO_3_, 1.25 mM NaH2PO4, 1 mM MgSO4, 10 mM D-glucose saturated with 95% O_2_/5% CO_2_, pH 7.5, PCO_2_ 35 mmHg.

### Imaging assay of gap junction transfer

2-Deoxy-2-[(7-nitro-2,1,3-benzoxadiazol-4-yl)amino]-D-glucose, NBDG, was included at 200 μM in the patch recording fluid, which contained: K-gluconate 130 mM; KCl 10 mM; EGTA 5 mM; CaCl_2_ 2 mM, HEPES 10 mM, pH was adjusted to 7.3 with KOH and a resulting final osmolarity of 295 mOsm. Cells were imaged on a Cleverscope (MCI Neuroscience) with a Photometrics Prime camera under the control of Micromanager 1.4 software. LED illumination (Cairn Research) and an image splitter (Optosplit, Cairn Research) allowed simultaneous imaging of the mCherry tagged Cx26 subunits and the diffusion of the NBDG into and between cells. Coupled cells for intercellular dye transfer experiments were initially selected on the basis of tagged Cx26 protein expression and the presence of a gap junctional plaque, easily visible as a band of mCherry fluorescence (e.g. Figures 1 and 2). After establishing the whole cell mode of recording, images were collected every 10 seconds.

**Figure 1.**
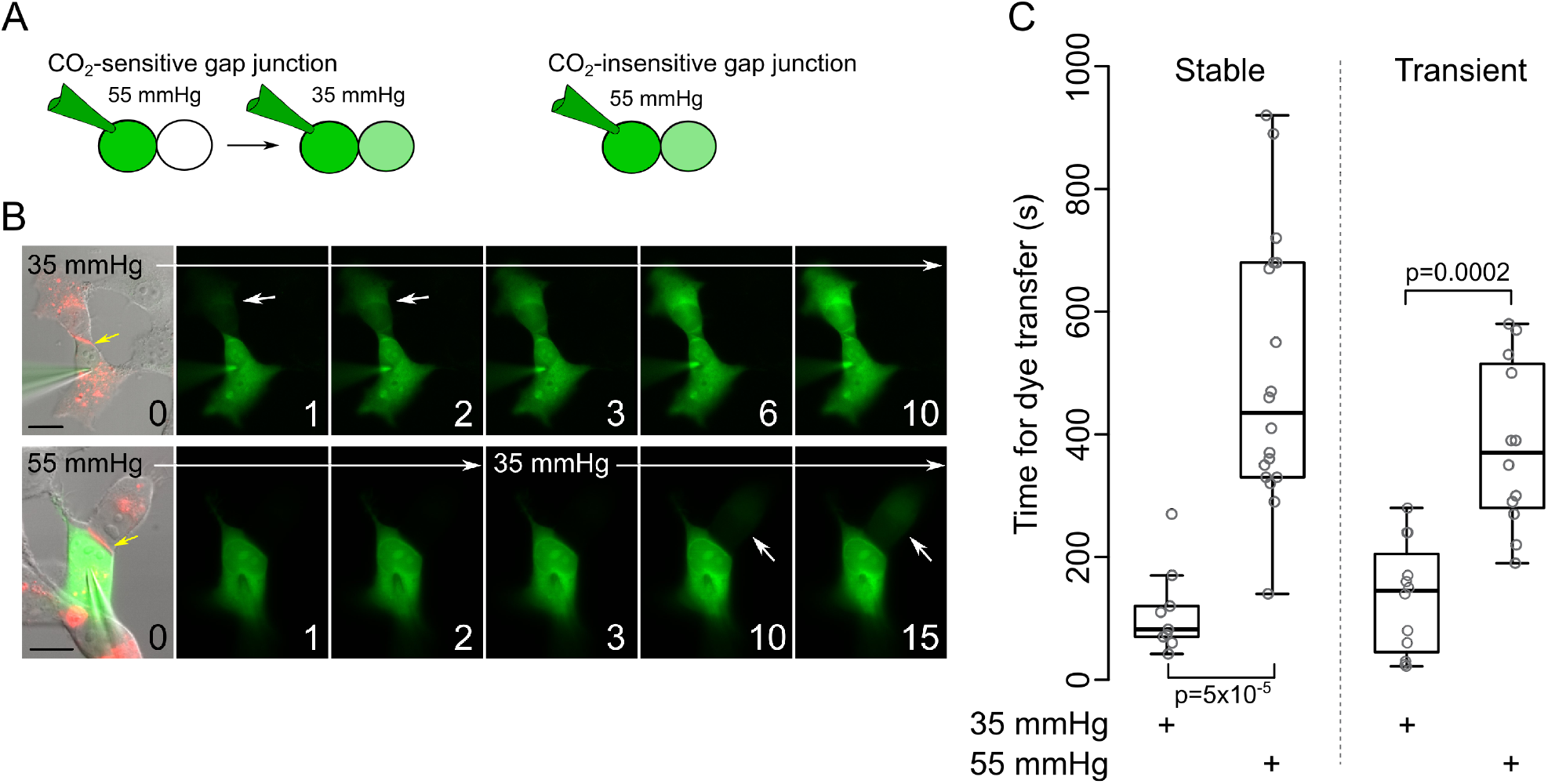
Dye transfer assay to assess the CO_2_ sensitivity of gap junction coupling. A) Logic of the assay: a CO_2_ sensitive gap junction will display no or very little dye transfer from the donor to acceptor cell under conditions of high PCO_2_. This will occur once the cells are transferred to a low PCO_2_ saline. A CO_2_ insensitive gap junction will exhibit equally rapid dye transfer at low and high PCO_2_. B) Images showing dye transfer between coupled cells at two different levels of PCO_2_. Numbers in lower right corner are recording time in minutes. Picture at 0 minutes in both rows is a merge of DIC and mCherry (for Cx26, red) and NBDG fluorescence (green) and is the beginning of the recording. The gap junction can be observed as a red stripe between the coupled cells (yellow arrows, both rows). Subsequent pictures show just the NBDG fluorescence. In 35 mmHg PCO_2_ dye transfer to the acceptor cell is evident after 1 minute (white arrows). Starting the recording at a PCO_2_ of 55 mmHg, and then transferring to 35 mmHg saline after 2 minutes greatly slows dye transfer; fluorescence in the acceptor cell is only seen after 10 minutes (white arrows). Scale bar is 20 μm. C) Summary data for stably expressing (mouse Cx26, *n*=9 and 18 for low and high CO_2_ respectively) and transiently transfected (human Cx26. tagged with mCherry, *n*=12 for low and high CO_2_) HeLa cells, showing the time for dye in the acceptor cell to reach 10% of the donor starting in 35 mmHg, and starting in 55 mmHg but transferring to 35mmHg after two minutes. When the recordings are commenced in 55 mmHg the dyes transfer time is much longer. Statistical comparisons Mann Whitney U test.

**Figure 2.**
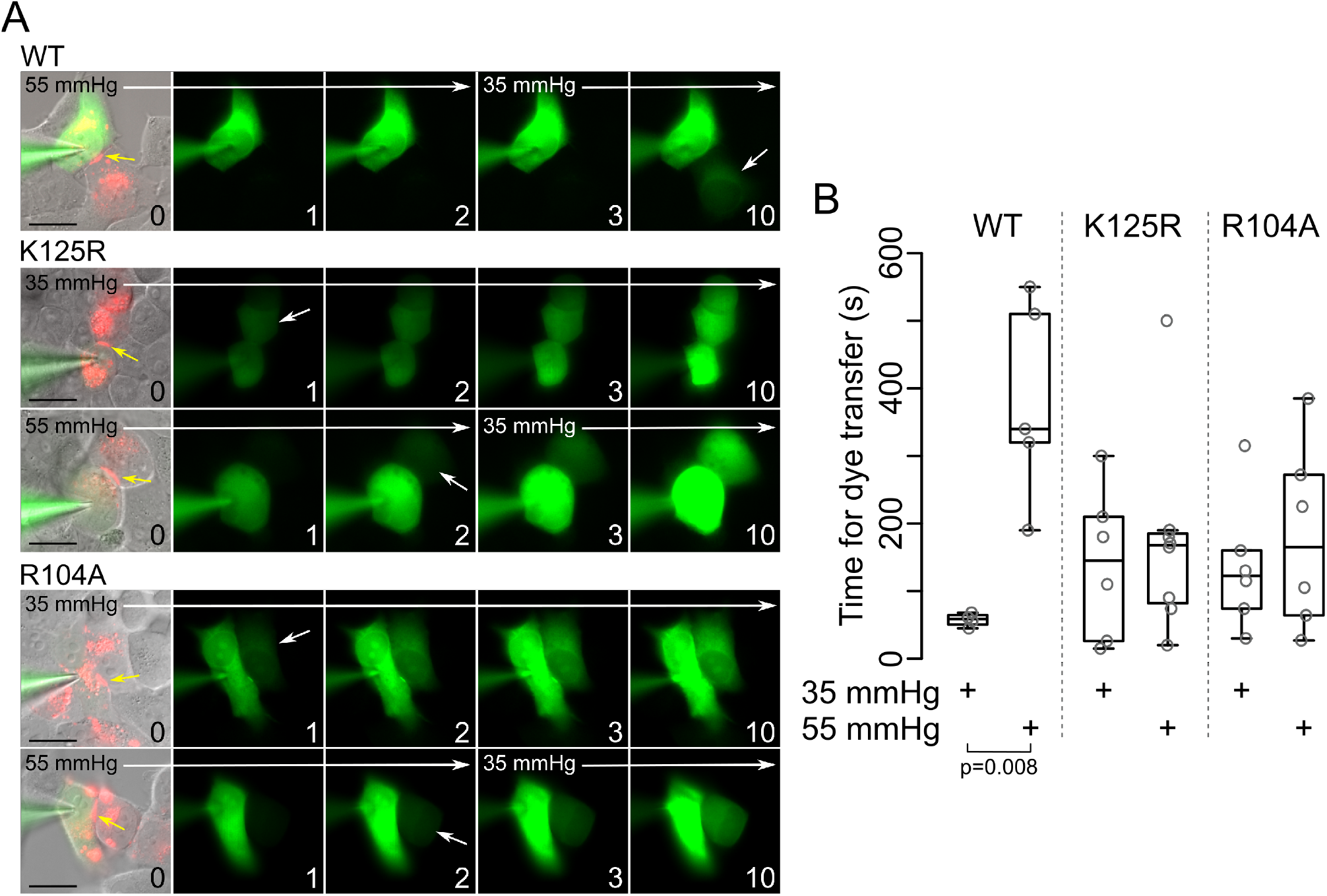
CO_2_ dependence of Cx26 gap junction closing depends on the carbamylation motif. A) Images showing permeation of NBDG through Cx26^WT^, Cx26^K125R^ or Cx26^R104A^ at different levels of PCO_2_. Unlike the Cx26^WT^, where elevated PCO_2_ slows permeation of NBDG to the coupled cell, permeation of NBDG between cells coupled by Cx26^K125R^ or Cx26^R104A^ and apparently unaffected by PCO_2_. White arrows indicate appearance of NBDG in the coupled cell, yellow arrows highlight the gap junction structure, scale bar represents 20 μm, numbers in bottom right corners indicate time of image in minutes after start of recording. B) Summary data shows that rat Cx26^WT^ gap junctions are CO_2_ sensitive (55 mmHg PCO_2_ delays the passage of dye across the gap junction), but those comprised of rat Cx26^K125R^ or rat Cx26^R104A^ show no CO_2_-dependence in the time required for dye transfer from the donor to acceptor cell. Mann Whitney U test WT 35mmHg vs WT 55 mmHg, *p*=0.008.

### Patch Clamp recordings from coupled cells

Cover slips containing non-confluent cells were placed into a perfusion chamber at room temperature in sterile filtered standard aCSF. Two Axopatch 200B amplifiers were used to make whole-cell recordings from pairs of HeLa cells. The intracellular fluid in the patch pipettes contained: K-gluconate 130 mM, KCl 10 mM, EGTA 10 mM, CaCl_2_ 2 mM, HEPES 10 mM, sterile filtered, pH adjusted to 7.3 with KOH. An agarose salt bridge was used to avoid solution changes altering the potential of the Ag/AgCl reference electrode. All whole-cell recordings were performed at a holding potential of −50 mV. Steps to −40 mV were applied to each cell in alternation to measure the whole cell and gap junction conductances. The current flow from the cell at −40 mV to the cell at −50 mV flows through the gap junction and can thus be used to calculate the gap junction conductance. The outward current during the +10 mV step in each cell represents the whole cell conductance which is a combination of all current pathways out of the cell. These comprise any intrinsic conductances and also the gap junction conductance itself.

### Elastic network modelling

Elastic network model (ENM) simulations were performed based on the regular implementation using PDB file 2ZW3, where all the Cα atoms in the protein within a given cut-off radius (8 Å) were joined with simple Hookean springs (Tirion, 1996; Rodgers *et al.*, 2013a). The spring constants were set to a constant value of 1 kcal mol^−1^ Å^−2^. The presence of CO_2_ was represented in the ENM by the inclusion of an additional Hookean spring between residues K125 and R104 of each set of neighbouring monomers, following the same procedure previously used for ligand binding (Rodgers *et al.*, 2013b). The mass-weighted second derivatives of this potential energy matrix were diagonalised to produce eigenvectors, *e*, which are the normal modes, and eigenvalues which are the squares of the associated frequencies, *ω*, and can be used to calculate the free energy of each mode.

The first six modes, that is the lowest frequency modes, represent the solid body translational and rotational motions of the protein and are thus excluded from the analysis. The overlap of the modes in the unbound and CO_2_ bound states were calculated by comparison of the eigenvectors (Rodgers *et al.*, 2013a). A value of 1 indicates that the motions are identical whereas a value of 0 indicates that the motions are completely different.

### Statistics and Reproducibility

Statistical analysis was performed with the R language. Data has been plotted as box and whisker plots, with individual points superimposed, where the box is interquartile range, bar is median, and whisker extends to most extreme data point that is no more than 1.5 times the interquartile range. Each individual point is from a single patch clamp experiment and is counted as a replicate. Statistical comparisons were performed with the Krusak Wallis test (for multiple comparisons) or Mann Whiney U test. Exact p values are shown for comparisons that showed a significant difference between samples.

## Results

### Rate of dye transfer between coupled cells shows CO_2_-dependence of Cx26 gap junctions

Many investigators have used dye transfer assays to demonstrate the presence of gap junctions (Spray *et al.*, 1991; Elfgang *et al.*, 1995; Abbaci *et al.*, 2007). We therefore adapted this type of assay to test whether increases in PCO_2_ could alter the permeation of a fluorescent glucose analogue (2-Deoxy-2-[(7-nitro-2,1,3-benzoxadiazol-4-yl)amino]-*D*-glucose, NBDG) through Cx26 gap junctions formed between HeLa cells. Starting with HeLa cells that stably expressed untagged Cx26, we made whole cell recordings from a single cell in a coupled pair. After introducing NBDG, via the patch pipette, into one of the cells (the *donor*) of a coupled pair, we recorded the time taken for the dye to diffuse into the coupled (*acceptor*) cell and achieve 10% of the fluorescence intensity of the donor cell (Figure 1A). If the gap junction is sensitive to CO_2_, then permeation of NBDG from the donor to acceptor cell should be altered by elevated PCO_2_ (Figure 1A). By recording from pairs of HeLa cells that stably expressed Cx26, we found that NBDG, rapidly permeated into coupled cells in solutions with a PCO_2_ of 35 mmHg (Figure 1C). When the recording was initiated in a saline with a PCO_2_ of 55 mmHg, there was little permeation during the period of high PCO_2_ and this only occurred once the PCO_2_ was reduced to 35 mmHg (Figure 1C). We refined this assay by performing it on cells that had been transfected with Cx26 tagged with mCherry. This allowed us to directly visualize the gap junctions and thus select pairs of coupled cells for the assay (Figure 1B). Once again, we found that the level of PCO_2_ altered the time it took for NBDG to permeate from the donor to the acceptor cell (Figure 1B, C).

Having established the validity of the fluorescence assay to detect CO_2_-sensitive intercellular coupling, we next investigated whether the CO_2_-sensitivity of Cx26 gap junctions depended on the same residues (K125 and R104) that are necessary for the CO_2_-sensitivity of the Cx26 hemichannel (Meigh *et al.*, 2013). We examined the effect of two mutations K125R and R104A that individually remove CO_2_ sensitivity from the hemichannel. Cx26^K125R^ and Cx26^R104A^ formed gap junctions between HeLa cells (Figure 2A) that were readily permeated by NBDG. Our data showed that unlike wild type Cx26 gap junctions, CO_2_ made no difference to the rate of dye transfer between cells coupled via Cx26^K125R^ or Cx26^R104A^ (Figure 2A, B). CO_2_-dependent gap junction closure appears to require the same residues that are essential for hemichannel opening.

### Electrical coupling via Cx26 gap junctions is sensitive to CO_2_

To confirm the observations made during intercellular dye transfer assays, we made simultaneous whole cell recordings from isolated pairs of Cx26-expressing HeLa cells that were in close apposition, as these cells had a high probability of being electrically coupled. On establishing whole cell recordings from each cell, the cells were clamped at a holding potential of −50 mV. We used a simple protocol of repeated 10 mV steps, alternating in each cell (see Methods). In all the illustrations, the downward currents during the voltage steps are proportional to the junctional conductance. Untransfected parental HeLa cells could exhibit gap junction coupling but this was insensitive to CO_2_ (median change in conductance −0.03 nS, 95% confidence limits −0.2 and 0.11, *n*=10; Figure 3A, B). In cells that stably expressed mouse Cx26, application of hypercapnic saline (PCO_2_ 55 mmHg) at a constant extracellular pH of 7.5 caused a reduction of junctional coupling (Figure 3A, B). In some cases, this was a partial effect and, in other cases, the coupling between cells was completely abolished. Overall the median change of the gap junction conductance (from control to 55 mmHg PCO_2_) was −5.6 nS (95% confidence limits −11.9 and −3.2 nS, *n*=29).

**Figure 3.**
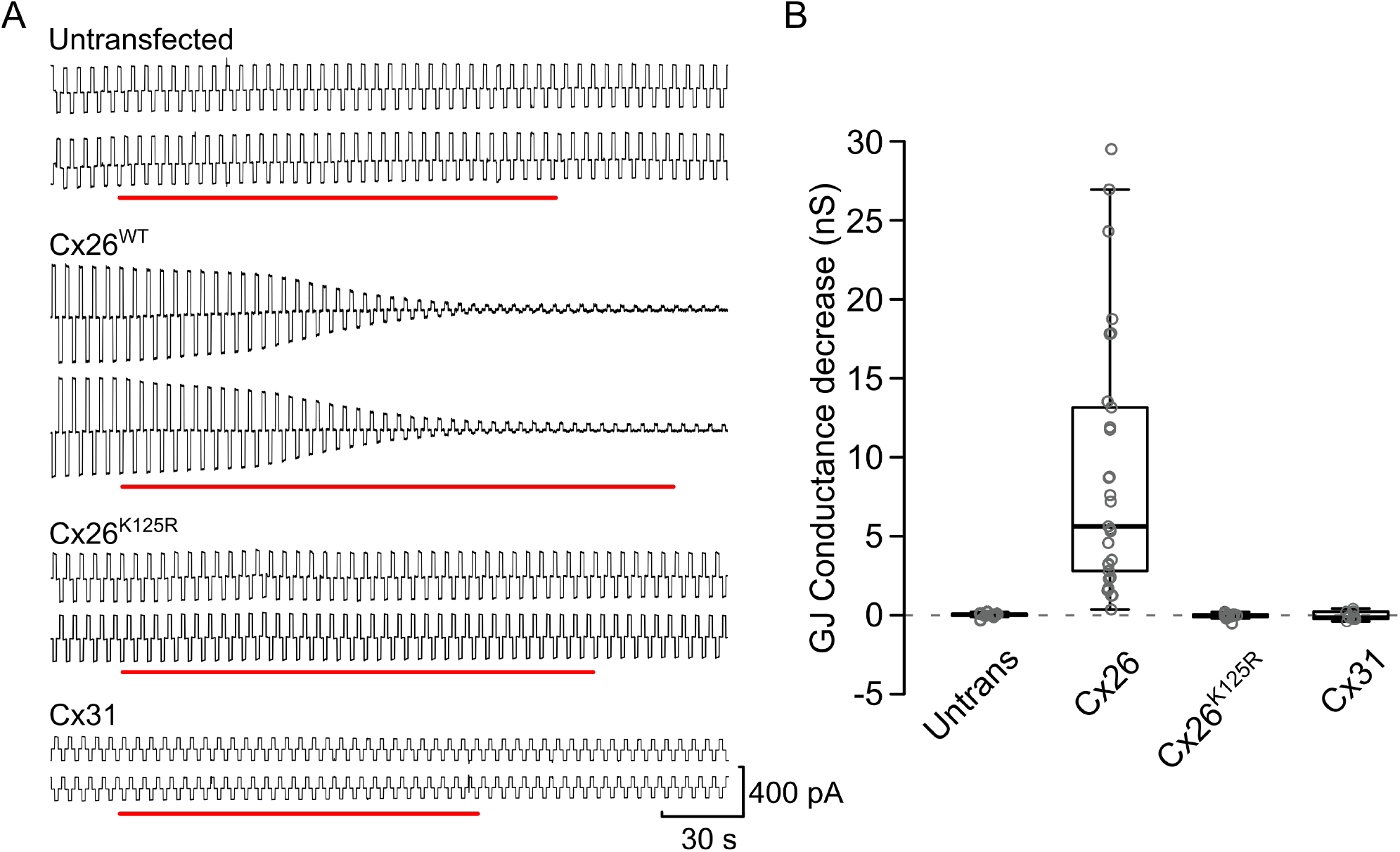
The CO_2_-dependent closure of gap junctions is specific to Cx26^WT^. A) Represenstative examples of electrophysiological recordings from cells displaying gap junction coupling. In each case, both cells were voltage clamped at −50 mV. On top of the holding potential, alternating +10 mV (1.25 s in duration) steps were applied to each cell. The upward currents represent the whole cell current, which will include the gap junction current; the downward currents represent current flow from the cell at −40 mV to the cell at −50 mV and is thus current flow through the gap junction. Untransfected HeLa cells can exhibit coupling, but this is not sensitive to CO_2_. Gap junctions formed between HeLa cells stably expressing mouse Cx26^WT^ are rapidly and completely blocked by CO_2_. Gap junctions formed by rat Cx26^K125R^ are unaffected by CO_2_, as are rat Cx31 gap junctions. Red bars represent the application of saline with 55 mmHg PCO_2_. B) Summary data showing the effect of CO_2_ on gap junction coupling between untransfected (parental) HeLa cells, and HeLa cells expressing Cx26^WT^, Cx26^K125R^ or Cx31.

We next tested the requirement for the carbamylation motif for gap junction closure by CO_2_ in the electrophysiological assay. Gap junctions formed between HeLa cells expressing Cx26^K125R^ were insensitive to a PCO_2_ of 55 mmHg (Figures 3A, B; median change in conductance 0.01 nS, 95% confidence limits 0.2 and −0.06 nS, *n*=14). Furthermore, gap junctions of Cx31, which lacks the carbamylation motif (Meigh *et al.*, 2013), were also insensitive to a PCO_2_ of 55 mmHg (Figure 3A, B; median change in conductance 0.07 nS, 95% confidence limits 0.44 and −0.29 nS, *n*=11).

We have therefore used two independent methods to demonstrate that Cx26 gap junctions are closed by moderate changes in CO_2_ (a PCO_2_ of 55 mmHg) and this CO_2_-dependent closure depends upon the same residues (K125 and R104) that are required for hemichannel opening by CO_2_.

### CO_2_-dependent and pH-dependent closure of Cx26 gap junctions can be dissociated by the K125R mutation

A possible mechanism by which CO_2_ could close the gap junction is via intracellular acidification. CO_2_ is known to permeate membranes and, by combining with water, can acidify the intracellular milieu. In prior studies, very high levels of CO_2_ (30-100%) have been used as a pharmacological method to close a variety of gap junctions (Spray *et al.*, 1991; Young & Peracchia, 2004). We therefore tested whether intracellular acidification, imposed independently of CO_2_ by application of 30 mM propionate (Jahromi *et al.*, 2002; Haussig *et al.*, 2008), could close the Cx26 gap junction, and whether this could be altered by the K125R mutation, which eliminates CO_2_-dependent closure. This level of propionic acid will cause far greater intracellular acidification than that caused by the elevated CO_2_ used in this study (Cook *et al.*, 2012). Propionic acid treatment reduced the gap junction conductance of wild type Cx26 by 5.3 nS (median, 95% confidence limits 2.8 and 8.3 nS, *n*=12, Figure 4). This manipulation also reduced the gap junction conductance of Cx26^K125R^ by a similar amount 7.7 nS (median, 95% confidence limits 4.1 and 11.0 nS, *n*=13, Figure 4). The actions of CO_2_ and intracellular pH on the Cx26 gap junction conductance are thus of similar magnitude, but mechanistically independent. Specifically, pH-induced gap junction closure does not require K125, whereas CO_2_-dependent closure does require this residue. This in turn suggests that CO_2_ has a direct action on Cx26, most probably via the carbamylation reaction that we have proposed for the opening of hemichannels (Meigh *et al.*, 2013), to cause closing of the gap junction.

**Figure 4.**
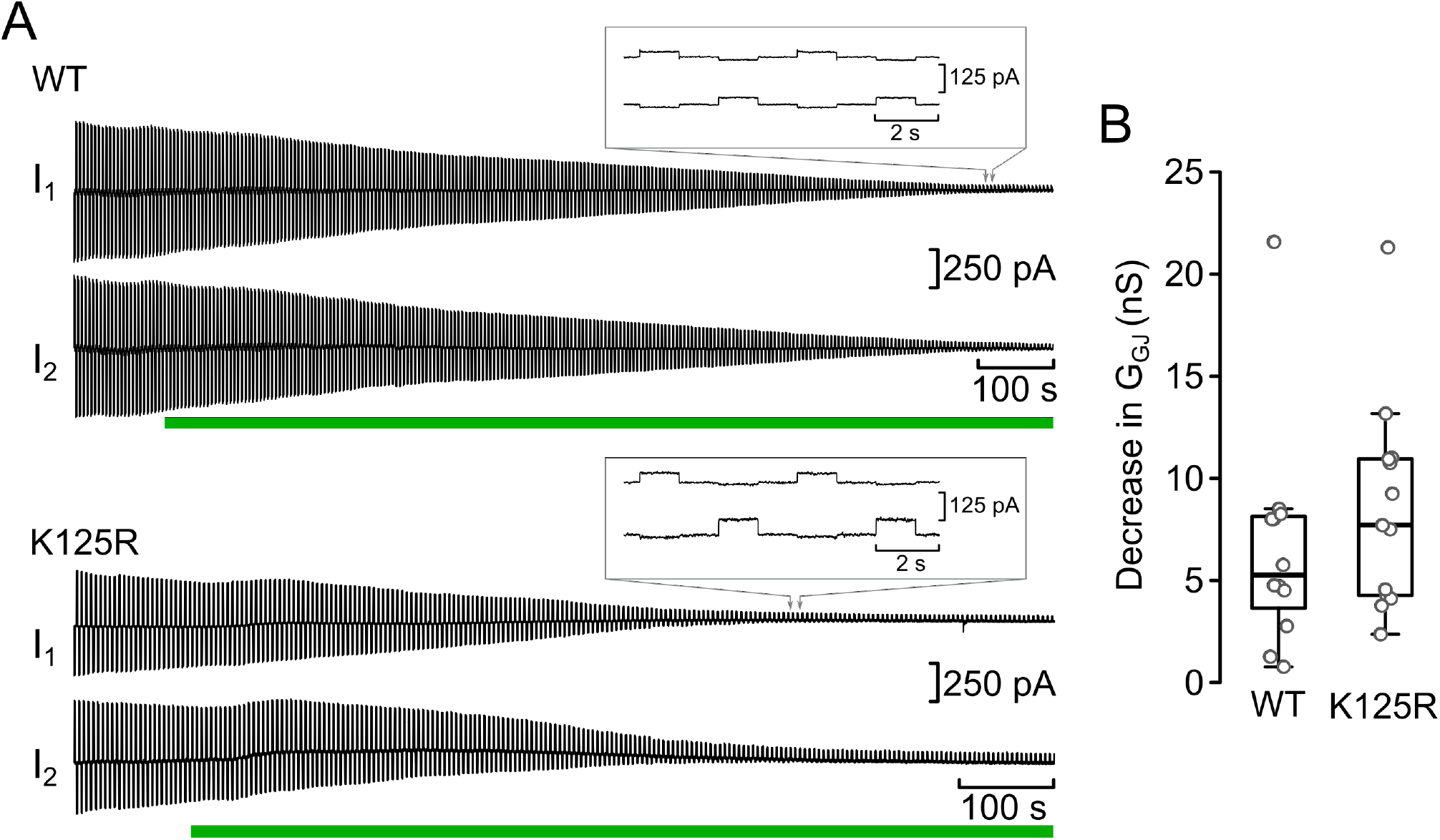
Intracellular acidification closes Cx26 gap junctions and is independent of the carbamylation motif. a) Sample recordings of gap junctions formed between cells transiently expressing human Cx26^WT^ or human Cx26^K125R^ tagged with mCherry. Alternating +10 mV steps were applied to each cell, the downward currents represent the flow of current through the gap junction to the coupled cell. Application of 30 mM propionate (green bar) caused a reduction in the gap junction currents, that was similar for Cx26^WT^ and Cx26^K125R^. The insets show the current traces at indicated time when Gap junction blockade was almost complete. b) Summary data showing the gap junction conductance decrease caused by acidification of HeLa cells expressing Cx26^WT^ or Cx26^K125R^.

### Effect of KID syndrome mutations on CO_2_-dependence of gap junction coupling

We have previously shown that KID syndrome mutations A88V, N14K, and A40V (Figure 5) abolish the ability of CO_2_ to open the mutant hemichannels (Meigh *et al.*, 2014; de Wolf *et al.*, 2016; Cook *et al.*, 2019). Recently, we discovered that certain KID syndrome mutations induce alternative splicing of the Cx26 mutation when this is tagged with a fluorescent protein (Cook *et al.*, 2019). This splicing results in poor expression and cell death but can be prevented by also mutating the 5’ splice site (Cook *et al.*, 2019). However, the 5’ splice site cannot be silently mutated so we chose a conservative mutation, M151L (Figure 5), which appeared to have little effect on Cx26 expression by itself. Cx26^M151L^ hemichannels are blocked normally by extracellular Ca^2+^ and they retain CO_2_-sensitivity (Cook *et al.*, 2019). Furthermore, Cx26^A40V,M151L^ hemichannels are insensitive to CO_2_ (Cook *et al.*, 2019).

**Figure 5.**
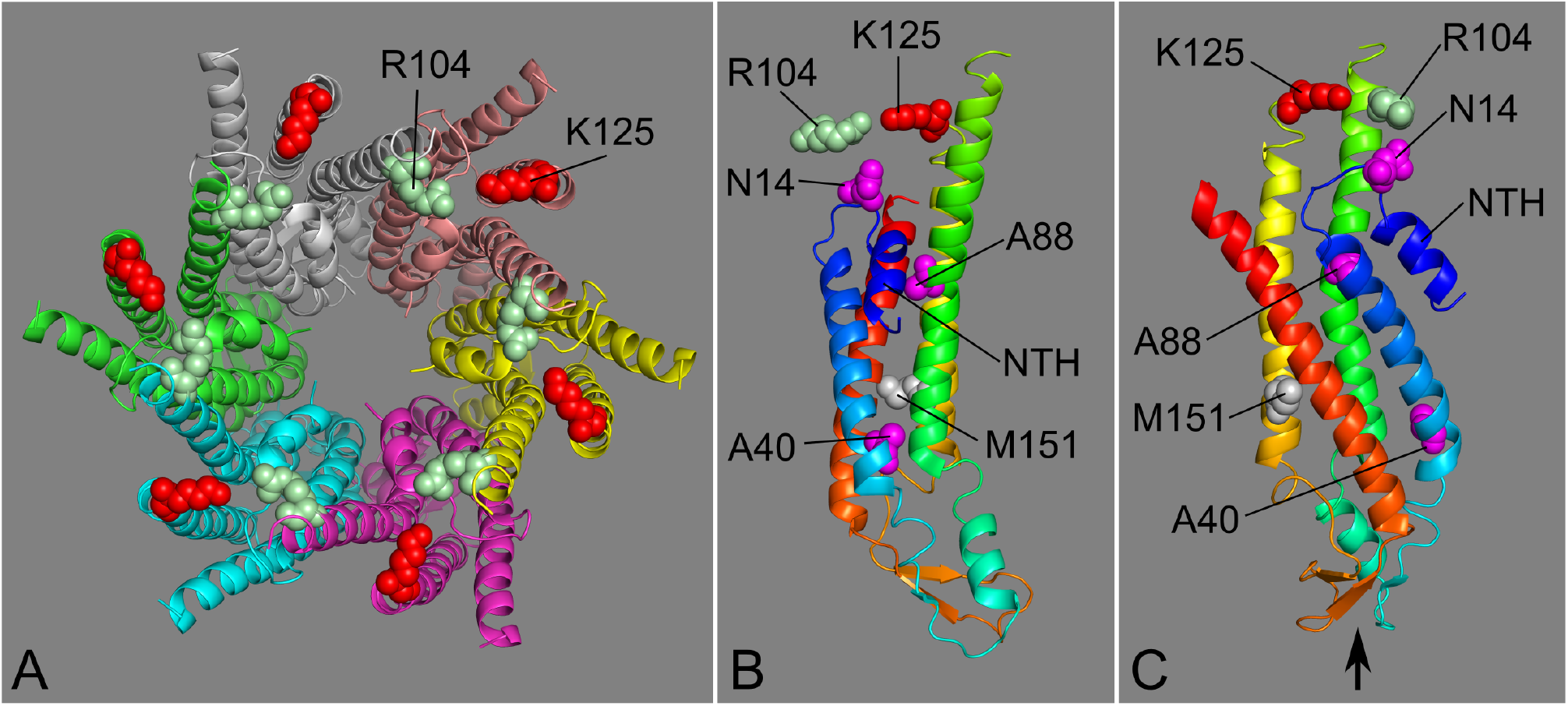
Position of the Cx26 carbamylation motif relative to the KIDS residues mutated in this study. a) View of the hemichannel as seen from the cytoplasmic face. Residues K125 and R104 of the carbamylation motif are shown in red and light green respectively. b) Side view of a single subunit showing the positions of mutated residues (N14, A40 and A88 are mutated in KIDS, M151 was mutated to prevent alternative splicing of KIDS mutated mRNA). R104 from the adjacent subunit is shown aligned to K125 of the subunit. NTH: N terminal helix. c) Additional view of the subunit after rotating it approximately 90° counterclockwise. Arrow indicates the loops in the subunit that dock to a hexamer in an opposed membrane to form a gap junction. Structure based on 2zw3 from the Protein Database.

When the M151L mutation was combined with the KIDS mutations, A40V, A88V or N14K (Figure 5), in each case, the doubly mutated Cx26 was invariably capable of forming functional gap junctions through which NBDG could permeate (Figure 6). For all three mutations, the Cx26 gap junction remained open in the presence of 55 mmHg PCO_2_ (Figure 6), a concentration that would normally close the wild type gap junction (Figures 1–3). Thus, the KIDS mutations not only prevent CO_2_-dependent opening of the hemichannel, but also prevent CO_2_-dependent closing of the gap junction, at least when combined with M151L. Interestingly, the effect of N14K on hemichannel sensitivity to CO_2_ is less than the other KIDS mutations (de Wolf *et al.*, 2016) and this parallels the trend in our data that permeation of NBDG through the Cx26^N14K,M151L^ gap junctions, while still occurring, may be slightly slowed at the higher level of PCO_2_ (Figure 6C, D).

**Figure 6.**
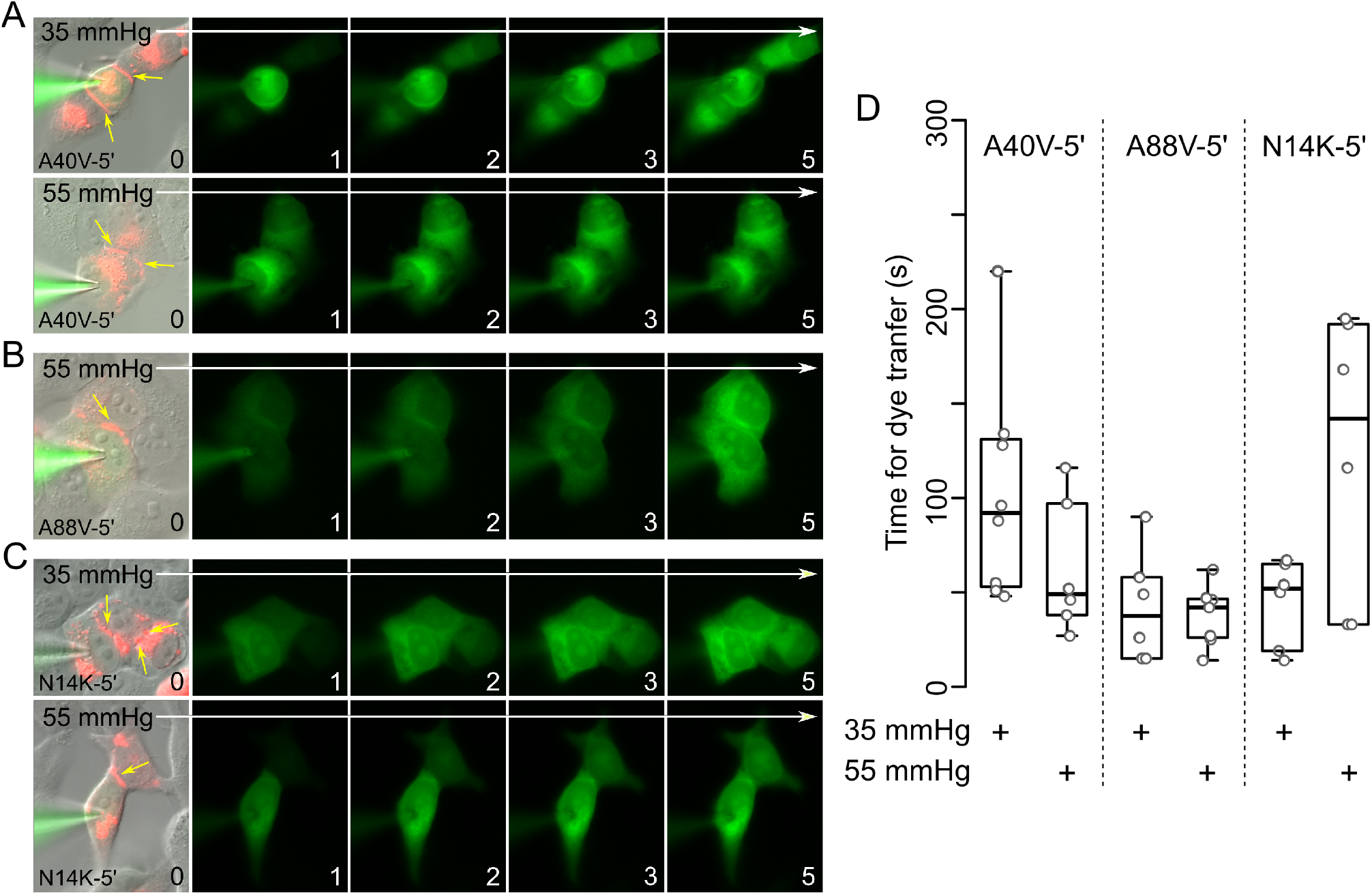
KIDS mutations of human Cx26 alter the CO_2_-sensitivity of Cx26 gap junctions. A) Gap junctions (indicated by yellow arrows) formed by Cx26^A40V-5’^ (combining the A40V mutation with M151L, to eliminate alternative splicing) are highly permeable to NBDG and lack any CO_2_ sensitivity. The images encompass 5 minutes of recording at a PCO_2_ of 35 mmHg (top) and 55 mmHg (bottom). Permeation of NBDG to the coupled cells happens within 2 minutes in both conditions. B) Gap junctions (indicated by yellow arrows) formed by Cx26^A88V-5’^ are insensitive to PCO_2_-rapid permeation occurs to the coupled cell even at a PCO_2_ of 55 mmHg. C) Gap junctions formed by Cx26^N14K-5’^ are not closed by CO_2_. Permeation of NBDG to coupled cells is shown at PCO_2_ of 35 mmHg (top row) and 55 mmHg (bottom row). D) Summary data showing the time for dye transfer to the coupled cell. While A40V-5’ and A88V-5’ show no difference in permeation time with PCO_2_, there is a tendency for the dye to permeate more slowly to the coupled cells for N14K-5’.

As the mutation M151L is a rare allele associated with non-syndromic hearing loss (Siemering *et al.*, 2006), we tested whether this mutation by itself affected gap junctions. Cx26^M151L^ permitted gap junction formation as gap junction plaques were clearly visible between expressing cells (Figure 7A). However, in 47 different recordings we did not observe either dye coupling (Figure 7B) or the electrophysiological hallmarks of electrical coupling (slowed capacitive charging currents during a voltage step indicative of current flow through the gap junction). By contrast, cells expressing Cx26^A40V,M151L^, Cx26^A88V,M151L^ and Cx26^N14K,M151L^ exhibited dye coupling in 14/15, 12/12 and 12/12 recordings respectively. These frequencies of coupling are statistically different χ^2^ = 82.2 (3 degrees of freedom, *p*=5.1 × 10^−18^). We conclude that the mutation M151L prevents permeation through gap junctions that are apparently formed but either remain closed at the transjunctional potential studied here, or are otherwise non-functional. This finding implies that the KIDS mutations appear to compensate for the deficiency in gap junction permeability introduced by the M151L mutation.

**Figure 7.**
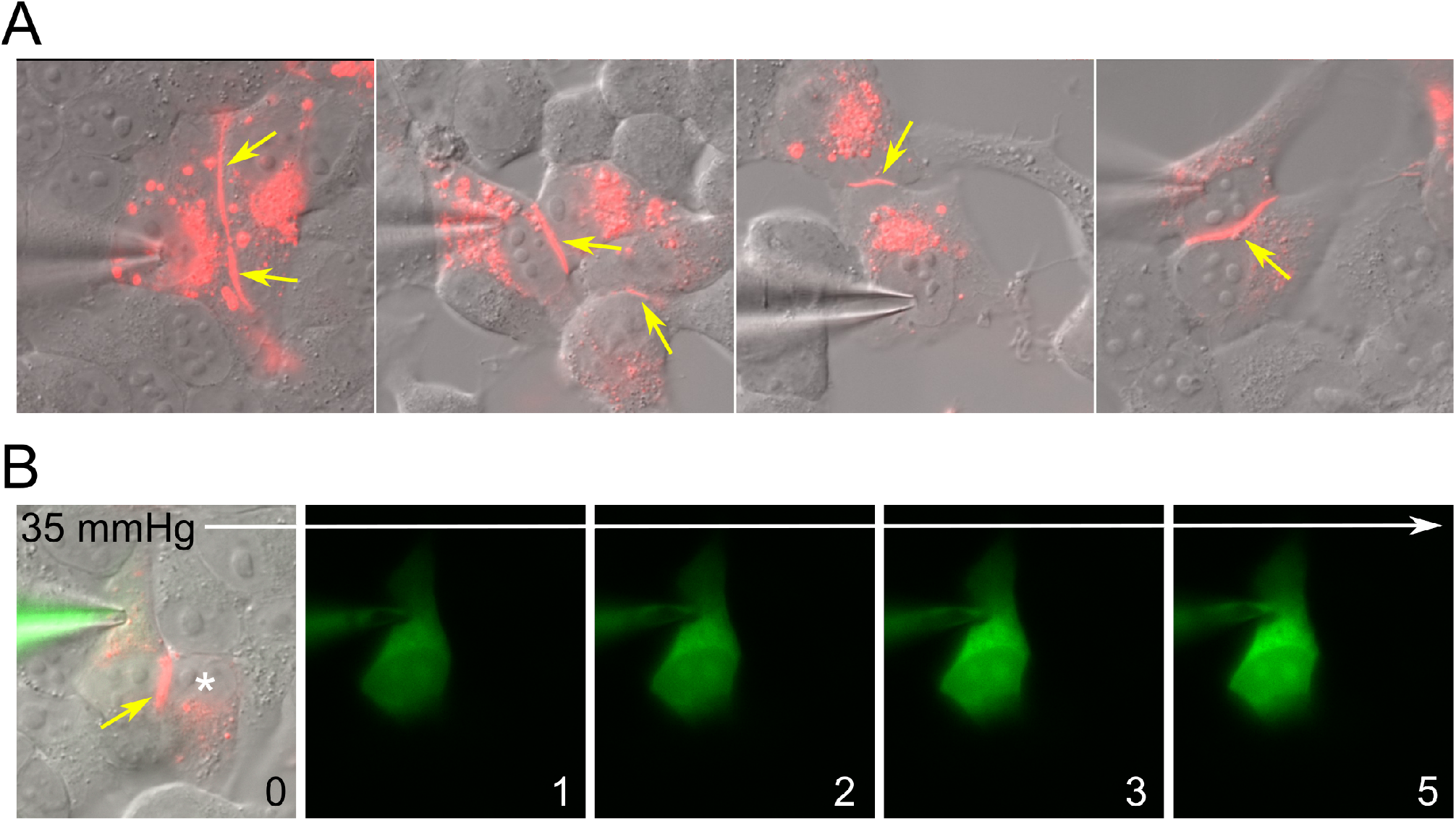
Physical gap junction plaques form between HeLa cells expressing human Cx26^M151L^ tagged with mCherry, but are non-functional. A) The gap junction plaques between cells are indicated by the yellow arrows. For clarity, the green channel (NBDG) has not been merged with the mCherry and DIC images. B) Images of a cell loaded with NBDG that formed a gap junction plaque (yellow arrow) with a neighbouring cell (*). However, NBDG does not permeate to the neighbouring cell within 5 minutes even at a permissive level of PCO_2_ (35 mmHg). Numbers represent minutes from start of recording.

### Elastic network modelling

Previous experimental data points to the importance of carbamylation of K125 and the formation of a salt bridge to R104 in the adjacent subunit to facilitate Cx26 hemichannel opening in response to CO_2_. It is not clear from this mechanism alone why the dodecameric Cx26 gap junction would be closed by CO_2_. Previous work has used coarse-grained modelling to demonstrate a mechanism whereby CO_2_ constrains the Cx26 hemichannel in the open state (Meigh *et al.*, 2013). We therefore used further coarse-grained modelling to probe the difference in behaviour of the Cx26 hemichannel and gap junction. Coarse-grained modelling reduces protein atomistic complexity for efficient computational studies of harmonic protein dynamics and is particularly suited to examining the contribution of entropy to channel opening over millisecond time scales (Sherwood *et al.*, 2008). While it is not possible to be certain that such calculated dynamics are true in the absence of experimentally determined structures, coarse-grained modelling has helped to support and explain structural data for membrane protein conformational changes (Shrivastava & Bahar, 2006; Sherwood *et al.*, 2008; Zheng & Auerbach, 2011; Isin *et al.*, 2012).

In an Elastic Network Model (ENM) the Cα-atom coordinates of an atomic resolution structure are used to represent a protein structure. Global protein harmonic motions within the ENM consists of a defined number of modes, each of a characteristic frequency and representing a particular harmonic motion within the protein. ENMs reproduce protein global low frequency modes well in comparison to experimental data (Delarue & Sanejouand, 2002; Valadie *et al.*, 2003). We therefore built coarse-grained ENMs (Tirion, 1996) to gain insight into the mechanism by which CO_2_ maintains the Cx26 hemichannel in the open state but the Cx26 gap junction in the closed state. ENMs were constructed using the coordinates from high-resolution crystal structures for the Cx26 hemichannel and gap junction in the CO_2_ unbound state. CO_2_ was represented in the ENMs by the inclusion of additional Hookean springs between residues K125 and R104 of neighbouring monomers in both the hemichannel and the gap junction (Meigh *et al.*, 2013).

We examined the similarities between eigenvectors for the Cx26 hemichannel and gap junction to understand the changes in harmonic protein motion caused by CO_2_-binding. The main open-close mode in the Cx26 hemichannel is defined as the lowest frequency mode that ignores the solid body translational and rotational motions. The solid body translational and rotational motions consist of six modes and so the open-close mode is mode 7 (Meigh *et al.*, 2013). This open-close mode in the hemichannel in the absence of CO_2_ (mode 7) becomes reordered as mode 15 in the presence of CO_2_ (Figure 8A; red-bordered square). The main open-close mode in the gap junction in the absence of CO_2_ (mode 10; modes 7-9 represent motions between the two hexamer rings) becomes reordered as mode 24 in the presence of CO_2_ (Figure 8B; green-bordered square). Analysis of the overlap between the main open-close mode in the gap junction and hemichannel in the absence of CO_2_ reveals an almost complete overlap between mode of the hemichannel (mode 7) and the gap junction (mode 10) (Figure 8C, blue-bordered square). The basic open-close dynamics are therefore likely similar between gap junction and hemichannel.

**Figure 8.**
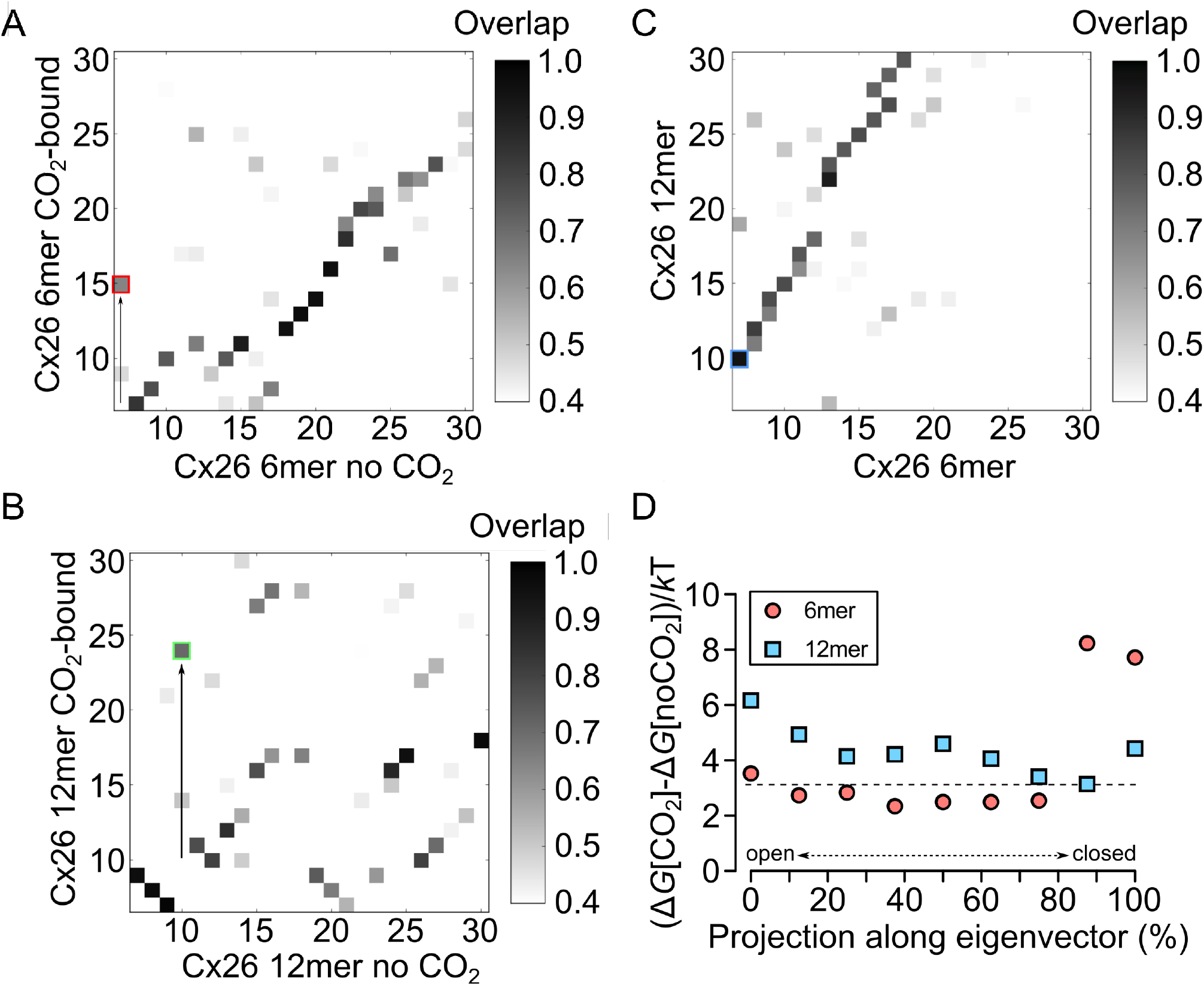
Analysis of the effect of CO_2_ on Cx26 hemichannel and gap junction gating via elastic network modelling. a) The main open-close mode in the hemichannel (6mer) in the absence of CO_2_ (mode 7) is reordered to mode 15 (red square) in the presence of CO_2_. Therefore, this mode contributes less to the total motion of the molecule in the presence of CO_2_-the hemichannel spends more time open (Meigh *et al.*, 2013). b) The main open-close mode in the gap junction (12mer) in the absence of CO_2_ (mode 10) is reordered to mode 24 (green square) in the presence of CO_2_. CO_2_ thus reduces the contribution of this mode to the total motion of the molecule, but it is not possible to state whether the gap junction is predominantly in the closed or open state from this analysis alone. c) Correspondence of the modes for the hemichannel (6 mer) and gap junction (12 mer). Mode 7 in the hemichannel corresponds very closely to mode 10 in the gap junction (indicated by blue square). d) For a range of partially open and closed eigenvectors for both the hemichannel and the gap junction, we then calculated the change in free energy on CO_2_-binding and thus the influence of CO_2_ on the open/closed eigenvector. The closer the free energy value is to zero, the less energy needed to bind CO_2_. The *x*-axis represents trajectory along the open/closed eigenvector going from fully open to fully closed. The *y*-axis represents the free energy for binding CO_2_ where the higher the value the less preferable is CO_2_-binding. For the hemichannel, CO_2_-binding is less energetically favourable in the closed state, whereas for the gap junction CO_2_-binding is favoured in the closed state.

To understand the differing roles of CO_2_ in the hemichannel and gap junction, we calculated a range of partially open and closed eigenvectors for both the hemichannel and the gap junction. We then calculated at each state, for both the hemichannel and gap junction, the free energy for CO_2_-binding and the influence of CO_2_ on the open/close eigenvector. This calculation provides a free energy for each state examined along the open/close eigenvector. Calculation of the free energy difference between the CO_2_-bound and unbound state provides information on the stability (binding energy of CO_2_ needed) for each state. In this case, the closer the value is to zero, the less energy needed to bind CO_2_ (Figure 8D). The *x*-axis of the Figure represents trajectory along the open/close eigenvector where the higher the value the more closed the hemichannel or gap junction. The *y*-axis of the Figure represents the difference in free energy for the CO_2_-bound and non-bound state where the higher the value is hypothesised to be less preferable for CO_2_-binding. On binding CO_2_, the hemichannel (6mer) or gap junction (12mer) will progress along the open/close eigenvector to make the difference in free energy more favourable for the CO_2_-bound state.

For the hemichannel, it is energetically favourable to bind CO_2_ in the open-state and then energetically unfavourable to close. For the gap junction it is energetically favourable to bind CO_2_ in the closed state and then energetically unfavourable to open. The differences in the open/closed state of the gap junction and hemichannel in response to CO_2_ can therefore be explained through entropy contributions to free energy change.

## Discussion

The key result from our study is that Cx26 gap junctions are closed by a direct action of CO_2_ on the protein. Our prior publications have demonstrated the opening action of CO_2_ on Cx26 hemichannels (Huckstepp *et al.*, 2010a; Meigh *et al.*, 2013; Meigh *et al.*, 2014; de Wolf *et al.*, 2016; de Wolf *et al.*, 2017; Cook *et al.*, 2019; Dospinescu *et al.*, 2019). This paper further shows that these diametrically opposite actions of CO_2_ on gap junctions and hemichannels depend on the same residues and presumably the same carbamate bridging mechanism.

Opposing modulation of gap junctions and hemichannels has been reported before. LPS and bFGF inhibit Cx43 gap junctions and open Cx43 hemichannels. While apparently similar to the results reported in this study, the modulation of these two entities is downstream of kinase activity and the signaling actions of arachidonate metabolites; i.e. these are indirect effects on Cx43 and are unlikely to involve the same residues in the protein (De Vuyst *et al.*, 2007). Similarly, the cytokines IL1β and TNFα also inhibit Cx43 gap junctions and open Cx43 hemichannels (Retamal *et al.*, 2007a). Both of these effects are mediated through a p38 MAPK dependent pathway. However, hemichannel opening triggered by these cytokines is sensitive to inhibition of nitric oxide synthase or changing redox state (Retamal *et al.*, 2007b), whereas the gap junction closure is insensitive to redox state. Thus, for the two entities, the mechanisms of modulation evoked by the p38 MAPK pathway differ (Retamal *et al.*, 2007a).

### Independence of pH- and CO_2_-dependent modulation of Cx26 gap junctions

A possible alternative interpretation of our data is that CO_2_ caused intracellular acidification and thus closed the gap junction. Under this interpretation, CO_2_ would have no direct action on Cx26. This interpretation is unlikely for two reasons. Firstly, only modest increases in PCO_2_ around the physiological norm were required to close Cx26 gap junctions. Such changes will only cause modest intracellular acidification. By contrast very profound acidification to pH values below 6.5 is required to close the Cx26 gap junction channel (Khan *et al.*, 2020). Secondly, and more importantly, the mutation K125R prevents the CO_2_-dependent closure of the gap junction, but does not affect the closing effect of acidification induced by application of propionate (Figure 4). The action of CO_2_ and pH on the gap junction are therefore mechanistically independent at the moderate levels of CO_2_ used in this study. This independence of mechanistic action of pH and CO_2_ is also true for Cx26 hemichannels as acidification causes hemichannel closure (Yu *et al.*, 2007), whereas a modest increase in PCO_2_ causes hemichannel opening (Huckstepp *et al.*, 2010a; Meigh *et al.*, 2013; de Wolf *et al.*, 2017; Cook *et al.*, 2019; Hill *et al.*, 2020).

Although the carbamylation motif in Cx26 is required for both the CO_2_-dependent opening of hemichannels and the CO_2_-dependent closure of gap junctions, the link between the CO_2_-dependent modulation of hemichannels and gap junctions is not immutable. Cx26 hemichannels of the lung fish *Lepidosiren* are insensitive to CO_2_ owing to the presence of an extended C-terminus, yet their gap junctions are still closed by CO_2_ (Dospinescu *et al.*, 2019). The contrary case is demonstrated by Cx32: hemichannels of this connexin can be opened by CO_2_, but Cx32 gap junctions are insensitive to the same doses of CO_2_ (Dospinescu *et al.*, 2019).

### Implications for structural biology of Cx26

The crystal structure of Cx26 shows the molecule in the form of a gap junction (Maeda *et al.*, 2009). The two connexons dock via interactions involving the two extracellular loops (E1 and E2) of each subunit. There are multiple hydrogen bonds formed between the opposing E1 and E2 loops of each connexon (Maeda *et al.*, 2009). These give a tight interaction that is likely to alter and constrain the conformation of hemichannels docked in a gap junction versus undocked hemichannels. The opposing modulation of gap junctions and hemichannels by CO_2_ must presumably arise from the conformational differences between free versus docked hemichannels.

Our ENM calculations of the interaction of CO_2_ with Cx26 hemichannels used the docked hemichannel from the gap junction as a model for the structure of the free hemichannel (Meigh *et al.*, 2013). This model allowed us to propose a plausible gating mechanism for CO_2_-dependent opening of the hemichannel, which we have supported with extensive mutational analysis (Meigh *et al.*, 2013). We exploited this knowledge to introduce novel gating mechanisms via the same intersubunit interactions by mutating Lys125 to Cys (Meigh *et al.*, 2015). This made the hemichannel NO-sensitive, via nitrosylation of the Cys125 and interaction with Arg104 of the neighbouring subunit. The double mutation −K125C, R104C - made the hemichannel redox sensitive (via redox modulation of disulfide bridges between subunits) (Meigh *et al.*, 2015). The mutated residues in these Cx26 variants are some considerable distance apart –far further than the short distances normally required for the ionic and covalent interactions between the respective residues. That these mutations were nevertheless effective at changing the gating of Cx26 implies considerable flexibility of the hemichannel that can bring apparently distant residues close enough to interact (Meigh *et al.*, 2015).

Despite this success in using a half gap junction as a model for the free hemichannel, the results in this study suggest the need for considerable caution in any future structural modeling of hemichannels. Not only may the isolated hemichannel be far more flexible than a hemichannel that is part of a gap junction, but it may also have a substantially different conformation. New structures of Cx26 hemichannels and gap junctions are therefore needed to probe their conformational differences, and provide firm mechanistic understanding of the differential modulation of hemichannels and gap junctions by CO_2_.

### KIDS mutations and Cx26 gap junctions

There are relatively few reports of the effects of KIDS mutations on Cx26 gap junctions. Prior reports suggest that the A40V mutation prevents Cx26 gap junction formation (Montgomery *et al.*, 2004). Clearly our data suggest that, in HeLa cells and in combination with the mutation M151L at least, the A40V mutation does not prevent formation of functional gap junctions (Figure 6a). Our data suggests further that N14K and A88V KIDS mutations (once again in combination with M151L) do not prevent gap junction formation and that these gap junctions are highly permeable to NBDG.

KIDS mutations are thought to cause the syndrome through gain-of-function. Several lines of evidence suggest that KIDS mutated Cx26 hemichannels are leaky: either they are less sensitive to Ca^2+^ blockade (Sanchez *et al.*, 2010; Mese *et al.*, 2011; Lopez *et al.*, 2013; Zhang & Hao, 2013; Sanchez *et al.*, 2014) or have an altered voltage sensitivity such that they spend more time in the open state at transmembrane potentials closer to the resting potential (Sanchez *et al.*, 2016; Valdez Capuccino *et al.*, 2019). Our finding that three KIDS mutations studied here prevent gap junctions from closing to CO_2_ is effectively a further gain of function caused by these mutations: they will remain open under conditions of elevated CO_2_. Interestingly, A40V hemichannels in oocytes are less sensitive to pH blockade than wild type Cx26 hemichannels, indicating that this mutation has multiple effects on the gating of these channels (Sanchez *et al.*, 2014). It is also interesting that the three KIDS mutations tested here could compensate for the loss of gap junction coupling induced by M151L. This observation is suggestive that these mutations enhance channel permeation and are also consistent with the gain-of-function hypothesis.

Although not a KIDS mutation, M151L has been reported to be associated with deafness (Siemering *et al.*, 2006). Our finding that it prevents functional gap junction communication suggests possible underlying mechanistic cause of hearing loss with this mutation. The endocochlear potential is essential for hearing - it provides the driving force for K^+^ to enter the hair cells through the mechanosensory ion channels present in their stereocilia. Cx26 gap junctions are a pathway for diffusion of K^+^ between the fibrocytes, basal cells and intermediate cells of the stria vascularis (Wangemann, 2006) and have been proposed to form part of the pathway that recycles K^+^ away from the hair cells and back into the endolymph, although this has been disputed (Zhao, 2017). Genetic deletion of Cx26 reduces the endocochlear potential by about half (Chen *et al.*, 2014). Given that M151L prevents gap junction coupling (although not the formation of gap junctions *per se*), we predict that it would reduce the endocochlear potential and that this is the reason for hearing impairment.

### Physiological implications

As Cx26 exists both as gap junctions and hemichannels, our findings have substantial physiological significance. In the context of breathing, Cx26 hemichannels are important and we have already explored their significance for chemosensory control (Huckstepp *et al.*, 2010b). However, there are reports of gap junctions in various nuclei implicated in the control of breathing (Dean *et al.*, 2002; Solomon *et al.*, 2003). CO_2_-dependent uncoupling of Cx26 gap junctions between cells may therefore be an additional mechanism that contributes to chemosensory control.

In lungfish and amphibia, hemichannels of Cx26 are insensitive to CO_2_ despite possessing the carbamylation motif (Dospinescu *et al.*, 2019). The extended C-terminal tail of Cx26 in these species interferes with hemichannel opening. However, the Cx26 gap junctions of these species can still be closed by CO_2_ (Dospinescu *et al.*, 2019). The ancestral function of the carbamylation motif in Cx26 was therefore most likely to close the gap junction and the new function of hemichannel opening only arose in the amniotes (Dospinescu *et al.*, 2019). Given that it has been conserved over many hundreds of millions of years, the CO_2_-dependent closure of Cx26 gap junctions must have some important physiological function. What this may be remains open to question, but we speculate that if a single cell in a coupled network were to be excessively metabolically active, it would act as a sink for metabolites such as ATP, glucose or lactate from the coupled cells. As the metabolically active cell would produce more CO_2_ this could be a self-regulating mechanism to preserve network integrity by uncoupling the “run-away” cell from the network thereby reducing its drain on the communal pool of metabolites. It is interesting in this context, that the PCO_2_ of renal cortex and liver (respectively 57 and 64 mmHg), two organs where Cx26 and Cx32 are abundantly expressed, is considerably higher than the PCO_2_ of systemic circulation (39-45 mmHg) (Hogg *et al.*, 1984).

## Acknowledgements

We thank the MRC (MR/P010393/1) (ND) and BBSRC (BB/S015132/1) (MC) for support. ND is a Royal Society Wolfson Research Merit Award Holder.

## Author contributions

SN, DM, ND: Acquisition and analysis of data

LM, EdW: Generation of mutants and stable cell lines

TR, MC: Elastic network modelling

ND wrote the paper, all authors commented on the final version.

## Data availability

All data generated or analysed during this study are included in this published article.

